# Comparison of biochemical, histological and morphological response of muscle-based condition indices to varying food levels in pre-flexion *Dicentrarchus labrax* (L.) larvae

**DOI:** 10.1101/023630

**Authors:** Ignacio A. Catalán, M. Pilar Olivar, Elisa Berdalet

## Abstract

The time-response, dynamics, classification power and relationship with survival of three muscle-based condition indices were analysed in pre-flexion sea bass (*Dicentrarchus labrax* L.) larvae in response to three feeding treatments: fed, non-fed and late-feeding. Larvae were reared at 19 ± 0.8°C and analysed from the second day of feeding (6 days after hatching (DAH)) to just prior the flexion stage (21 DAH). The indices were analysed in trunk muscle at biochemical (RNA/DNA, R/D), histological (percentage of muscle fibre separation, MFS) and morphometric (size-standardised body depth at the anus level, D_AZ_) level. No index was correlated with muscle length in the fed treatment regardless the period analysed (before or after full external feeding). The R/D was the only index that could detect past feeding differences in larvae ≤ 13 DAH. For larvae > 13 DAH, MFS, R/D and *D*_AZ_ in that order, yielded the best discrimination among treatments. For late-feeding larvae, latency was two days (R/D) and two-four days (MFS). Dynamics of recovery after food addition was also similar for R/D and MFS. Significant relationships between muscle growth and R/D or MFS were only found in larvae > 13 DAH. Both R/D and MFS encompassed the mass mortality event observed in non-fed larvae at 17 DAH. The *D*_AZ_ largely failed to give information on the feeding status.

## 1 Introduction

In the wild, the accumulated subtle action of growth-reducing factors (e.g. suboptimal feeding) can directly or indirectly explain most of the variation in larval mortality (Anderson 1988). Condition indices (CIs) are used to monitor physiological status or growth and operate at any organisation level of the individual. CIs differ in their sensitivity, latency, or dynamics (*sensu* Ferron and Leggett, 1994). One of the sources of variability of a given CI arises from tissular-specificity. The trunk (skeletal) muscle is a key tissue to study growth processes as it is catabolised during starvation and accounts for most on the mass increase in larvae of many species (e.g. Pedersen 1990). At a biochemical level, the RNA/DNA (R/D hereafter) measures the synthetic activity in the muscle of a growing larva, based on the more stable DNA content per cell with respect to RNA during metabolic processes (Buckley et al. 1999). At tissular level, suboptimal feeding decreases the thickness of the muscle fibres (e.g. Catalán and Olivar 2002) and muscle alteration is also reflected as a change in muscle shape (e.g. Koumoundouros et al. 2001). Size-independence is a must for CIs. This is usually accomplished by biochemical or histological indices, but not by morphometric indices. For these, the use of ratios, residuals or PCA (e.g. Suthers, 1992) has tried to remove the size effects, but have largely failed to do so due to the effects of allometry.

This work compares three size-independent muscle-based CIs at morphometric, histological and biochemical levels, in regard to i) their ability to differentiate feeding conditions of pre-flexion sea bass larvae and ii) encompass growth and mortality in the laboratory. We chose the sea bass *Dicentrarchus labrax* L. to compare these CIs, as the ontogeny, rearing requirements and growth of this species are known (Barnabé et al. 1976; Marangos 1986), as well as the effect of temperature and to a lesser extent food on muscle growth dynamics (Johnson and Katavic 1986; Bergeron and Person-Le Ruyet 1997; Ayala et al. 2003, López-Albors et al. 2003; Abdel et al. 2004; Alami-Durante et al. 2006; Catalán et al. 2007; López-Albors et al. 2008 and references cited therein).

## 2 Materials and Methods

Sea bass larvae from a single female of Mediterranean origin were reared at 19 ± 0.8°C since hatching in a small-scale recirculation system designed for experimental ecology. The rearing conditions, growth and survival were fully described in Olivar et al. (2000). Larvae were analysed from 6 to 21 days after hatching (DAH). The feeding treatments were: “fed” (*ad lib*), “non-fed” and “late-feeding” (starved individuals which were refed from 13 DAH (Table 1). Larvae were randomly sampled from 4l-rearing cylinders (described in Olivar et al. 2000), using 4 replicates (fed), 3 replicates (non-fed) and 1 replicate (late-feeding). Statistical analyses were performed using Minitab 15 (from Minitab Inc., U.S.A.). For parametric methods, homoscedasticity was always met.

**Table 1.** Number of larvae used for each index/day/feeding treatment. Late-feeding started at 13 days after hatching (DAH). The feeding scheme is in Olivar et al. (2000)

| Index | Treatment | Time (DAH) |  |  |  |  |  |  |  |  |  |  |  |  |  | Total *n* |
| --- | --- | --- | --- | --- | --- | --- | --- | --- | --- | --- | --- | --- | --- | --- | --- | --- |
|  |  | 6 | 8 | 10 | 11 | 12 | 13 | 14 | 15 | 16 | 17 | 18 | 19 | 20 | 21 |  |
| R/D | Fed | 5 | 6 | 5 | 4 | 4 | 5 | 4 | 6 | 3 | 8 | 6 |  |  |  | 56 |
|  | Non-fed | 6 | 5 | 6 | 4 | 2 | 3 | 4 | 2 | 1 | 3 | 6 |  |  |  | 42 |
|  | Late-feeding | - | - | - | - | - | - | 2 | 3 | 1 | 4 | 4 |  |  |  | 14 |
| MFS | Fed | 5 |  | 2 |  |  |  | 5 |  |  | 5 |  | 5 |  | 5 | 27 |
|  | Non-fed | 6 |  | 5 |  |  |  | 5 |  |  | 5 |  | 5 |  | 2 | 29 |
|  | Late-feeding | - | - | - | - | - | - | 4 |  |  | 5 |  | 3 |  | 4 | 16 |
| D<sub>AZ</sub> | Fed | 12 | 6 | 7 | 8 | 6 | 4 | 7 | 8 | 9 | 10 | 9 | 8 | 9 | 8 | 111 |
|  | Non-fed | 6 | 4 | 4 | 4 | 4 | 4 | 3 | 3 | 4 | 3 | 5 | 3 | 2 | 1 | 50 |
|  | Late-feeding | - | - | - | - | - | - | 4 | 3 | 2 | 4 | 4 | 3 | 3 | 4 | 27 |

### 2.1 Selection of size-independent condition indices

The CIs used embraced an expanded version (one more treatment) of previous biochemical and histological indices published by the authors (Catalán and Olivar 2002; Berdalet et al. 2005a), plus a new morphometric CI designed to remove the effects of size yet accounting for allometry. Larvae were sampled before being fed at 10 a.m. and stored at -80°C (for R/D) or in 10% phosphate-buffered formalin (rest of the analyses). Before processing, notochord length (*L*_N_), body depth at the anus level (*D*_A_) and head length (*L*_H_) (all in mm) were taken under a binocular microscope. Body measures were standardised to body muscle length (*L*_M_). The *L*_M_ was measured from the end of the chleitrum to the end of the notochord (*L*_N_-*L*_H_). All *L*_M_ measures were standardised to the length in the preserving formalin solution, using a conversion based on fresh larvae measured before and after placing them in that solution for 4 days (unpubl. data). The R/D was used as biochemical CI. Analyses were performed “mainly” on muscle tissue of individual defrozen larvae (heads and guts were discarded prior to processing) and analysed following Berdalet et al. (2005b).

The histological CI was published by Catalán and Olivar (2002) and is based on the semi-automatic detection of the percentage muscle fibre separation (MFS), applied to 3 μm saggital sections of Historesin (Leica)-embedded trunk muscle containing 4-6 myotomes and died with Lee’s Lee’s methylene-blue basic fuchin.

As morphometric CI, we selected the body depth at the anus level (*D*_A_). Size effects are not important if age is known (Fig. 1a) but it is not the case if age is unknown (Fig. 1b) as it happens in field samples. To account for allometric effects, the morphological factor of each individual (*D*_A_) was scaled to a common body size of the larval pool to be analysed (Lleonart et al. 2000). We divided the larvae into two groups to account for the two-phase growth observed in Fig. 2b. We selected the day at which no remnants of maternal reserved were observed (13 DAH) as a cut point. Allometry between *L*_M_ and *D*_A_ for each treatment and period was explored by fitting the equation

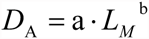

where b is the allometric factor, and a is the expected value of *D*_A_ at *L*_M_ = 1. The values a, b and the standard deviations (in ln-transformed units) were first calculated, and the existence of allometry tested through a T-test. If significant slopes did not differ from 1, the ratio *D*_A_/*L*_M_ was used, as ratios are uncorrelated with size in isometric growth. Otherwise, the *D*_A_ of each individual was normalised (*D*_AZ_) following Lleonart et al. (2000), according to

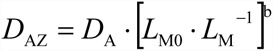

where *L*_M0_ is the selected reference *L*_M_ and b is the allometric parameter which relates *D*_A_ to *L*_M_. The slopes of the regressions were compared through GLM on ln-linearised variables (*L*_M_ used as a covariate).

**Fig. 1.**
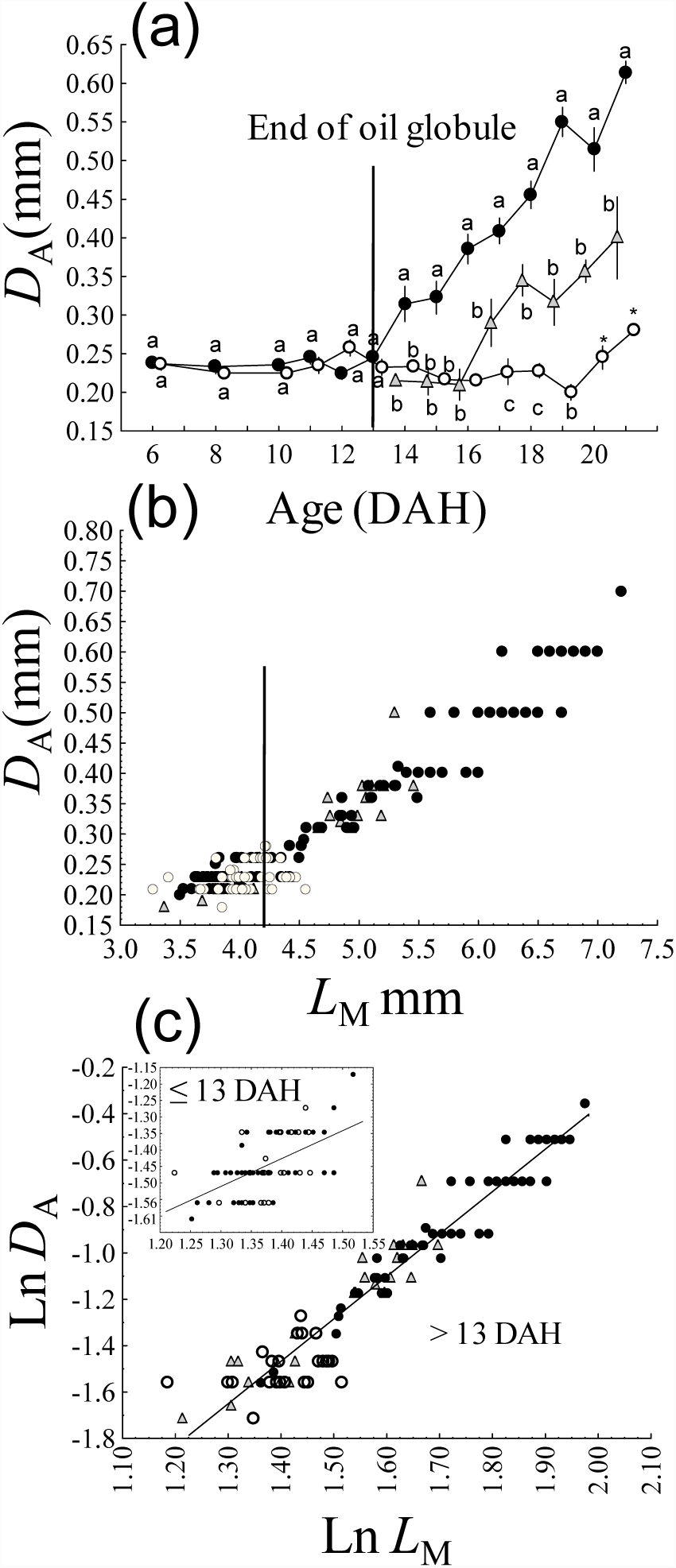
Evolution of body depth at anus level (D_A_) among feeding treatments *vs* age (a) and body muscle length (*L*_M_) (b). In (c), fitted regressions to both developmental groups are shown. For larvae ≤ 13 DAH: Ln *D*_A_ = 0.920 · Ln *L*_M_ - 2.711, F_1,67_, *r*^2^ = 0.33, *p* < 0.05. For larvae < 13 DAH: Ln *D*_A_ = 1.839 ·Ln *L*_M_ - 4.045, *F*_1,116_, *r*^2^ = 0.93, *p* < 0.001. Filled circles are fed treatment, open circles are non-fed treatment, triangles are late-feeding treatment. In (a), values are means ± 1 SE and, for a given day, significant differences among treatments are indicated by different letters (*p* < 0.05), using either t-tests (2 groups) or ANOVA (3 groups) with HSD for unequal number of observations.

**Fig. 2.**
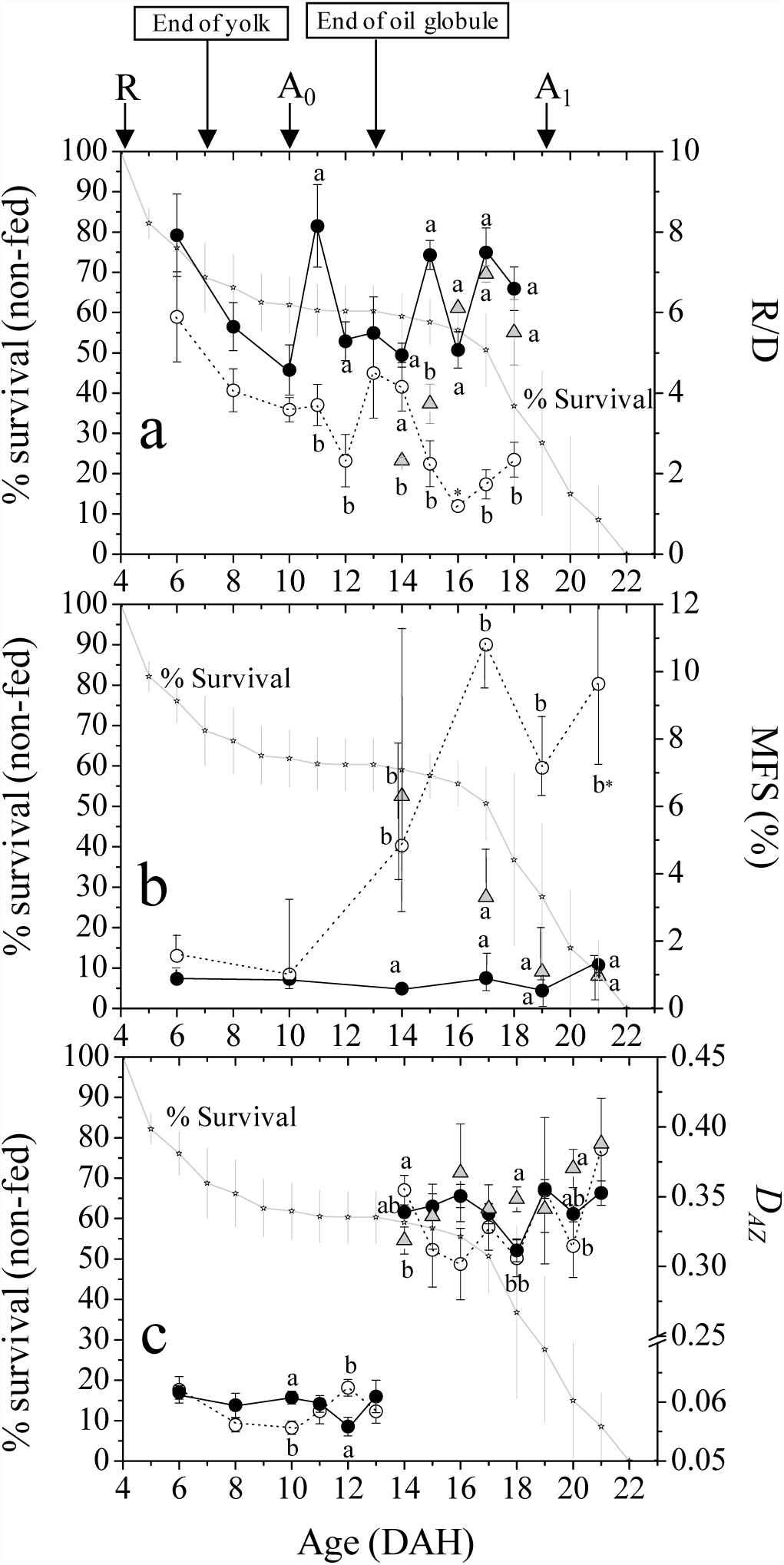
Relationship between percentage survival of non-fed larvae and the biochemical (a), histological (b) and morphometrical (c) condition indices for that treatment. Symbols, color codes and statistical tests as for Fig. 1, except for MFS, for which values are medians, errors are interquartilic ranges, and statistics are non-parametrical (Mann-Whitney for 2 groups and Kruskall-Wallis for 3 groups). The feeding scheme is indicated: R= rotifers, A_0_ = *Artemia* nauplii, A_1_= 1 day-old *Artemia* metanauplii.

### 2.2 Comparison of indices

The performance of each CI was explored with respect to age for each period (13 DAH cut point).

The *latency* (time to a detectable change in condition after a change in food supply) and *dynamics* (behaviour of this reaction) (Ferron and Leggett 1994) of each CI were analysed. Daily among-treatment differences were analysed through ANOVA followed by Tukey’s HSD in the case of R/D and *D*_AZ_, and through Kruskall-Wallis or Mann-Whitney tests for MFS due to the non-normality of the residuals. Their dynamics was modelled using standard regression analysis.

The “power” of each CI to assign a particular larva to its feeding treatment was assessed through linear Discriminant Analysis (DA). The DA was applied separately to the two developmental periods. As the sample size and sampling frequency differed for each index, two DA were applied: i) a conservative DA using coincident sampling days and sample size (scaling down the data by data point elimination to the number of data points of the CI with the lowest number of observations); ii) a second DA using all available data. The DA was performed setting *a priori* classification probabilities proportional to group sizes and with no cross-validation.

As no estimates of muscle dry weight was available, a volumetric proxy was used to estimate a daily muscle potential growth (*G*_MP_). For each day, it was measured as the expected absolute (*G*_MP_, mm^3^ day^-1^) or relative (% *G*_MP_) change in average muscle volume in the following day. The muscle volume was as assimilated to a cylinder according to

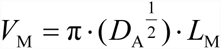

where *V*_M_ is muscle volume (mm^3^). *G*_MP_ was calculated as

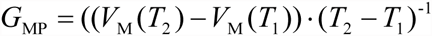

where *T*_1_ and *T*_2_ are time 1 and time 2 in days. The *G*_MP_ was used to estimate *V*_M_ at the following day (where gaps > 1 day appeared), and the % *G*_MP_ was estimated according to

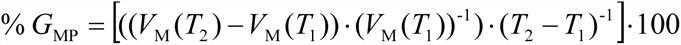

The average *V*_M_, *G*_MP_ and % *G*_MP_ were calculated for each day in the subsets of larvae in Table 1.

The percentage daily mortality (*M*, % day^-1^) was calculated as the average percent larvae dying with respect to the previous day. The correlation (Spearman, *r*_*s*_) between *G*_MP_, % *G*_MP_, *M* and the CIs was explored only in non-fed larvae (the only treatment showing a mass mortality event, Olivar et al. 2000). Data of larvae < 9 DAH were discarded because mortality was high and comparable across treatments before that day (see Olivar et al. 2000).

## 3 Results and Discussion

White fibres make the bulk of the preflexion muscle mass (Alami-Durante et al. 2006), and they are independent of small temperature differences during pre-flexion stages (Veggetti et al. 1990; Alami-Durante et al. 2006). Therefore, the skeletal muscle is an ideal tissue to study the effects of suboptimal feeding. Growth and survival was similar within replicated treatments (Olivar et al. 2000) and almost no larval mortality occurred in the fed larvae after 9 DAH. The major developmental events appear in Fig. 2.

### 3.1 Size-independence and choice of indices

The removal of size effects was successful. For any given feeding treatment, there were no significant differences in the *L*_M_ (ln-transformed) vs age relationships among groups of larvae used for the three CIs (GLM, age taken as covariate). For both R/D and MFS, no significant correlations with *L*_M_ were observed in fed or non-fed larvae in any period. However, a positive (R/D: *r*_*s*_ = 0.79, n = 14, *p* < 0.0001) and negative (MFS: *r*_*s*_ = -0.70, n = 24, *p* < 0.001) correlation was found in the late-feeding larvae. This is considered unavoidable in cases of sharp growth increase, and suggests that, in field studies, CIs should only be used in case of smooth growth functions. The DNA/RNA slope ratio of the standard curve was 3.965.

For the morphometric index, relationships between *D*_A_ and *L*_M_ were significant for the period ≤ 13 DAH, (fed: Ln *D*_A_ = 1.015 · Ln *L*_M_ - 2.837, *F*_1,41_, *p* < 0.005; non-fed: Ln *D*_A_= 0.698 · Ln *L*_M_ – 2.413, *F*_1,24_, p < 0.05). The lack of among-treatment significance in slopes or means (GLM) enabled pooling the groups for that first period. For larvae > 13 DAH, only the *D*_A_ of non-fed larvae was uncorrelated with *L*_M_. The remaining groups were significantly related (fed: Ln *D*_A_ = 1.827 · Ln *L*_M_ – 4.019, *F*_1,66_, *r*^2^ = 0.923, *p* < 0.05; non-fed: Ln *D*_A_ = 1.832· Ln *L*_M_ – 4.00, *F*_1,23_, *p* < 0.05). As no significant differences in slopes or means between fed and late-feeding larvae were found (GLM) and non-fed larvae had a very short size range (Fig. 1b), larvae were pooled for this period too (Fig. 1c). Slopes of the pooled groups differed between the two periods (GLM; slopes: *F*_1,184_, *p* < 0.05). Growth of *D*_A_ was isometric (H_0_: ß = 1, *p* = 0.56) in larvae ≤ 13 DAH, so ratios were derived to obtain *D*_AZ_. Larvae > 13 DAH showed a strong allometric development (Fig. 1c), hence the method by Lleonart et al. (2000) was applied to obtain D_AZ_, selecting a *L*_M0_ of 5 mm *L*_M_, which was the mean of the *L*_M_ distribution for all treatments. The resulting *D*_AZ_ values showed no correlation with *L*_M_ in the control group for any of the periods.

### 3.2 Comparison of indices properties

#### 3.2.1 Latency and dynamics

In Fed larvae, all CIs but R/D were stable through age and developmental period (Fig. 2). Non-fed larvae tended to show lower mean R/D values with respect to fed larvae (Fig. 2a). The MFS of non-fed larvae remained low until 10 DAH, increasing rapidly afterwards (Fig. 2b), which implies a similar latency to that of R/D (7-10 days). The concurrent decrease in mean R/D values in fed (-30%) and non-fed (-54%) larvae until 10 DAH was due to a higher increase rate in DNA content prior to that day (93% in fed, -1% in non-fed), which has been observed elsewhere (Bergeron and Person-Le Ruyet 1997), coupled to comparatively lower RNA figures (34 % in fed, -53% in non-fed). This might be related to high hyperplastic activity (Veggetti et al. 1990; Alami-Durante et al. 2006) occurring at the expense of maternal reserves. Data on *D*_AZ_ was difficult to interpret in terms of latency for the non-fed larvae (Fig. 2c). For late-feeding larvae, latency for mean R/D was 2 days, whereas for median MFS was 2-4 days (Fig. 2a,b). Latency values for mean *D*_AZ_ could not be computed (Fig. 2c).

The R/D dynamics of non-fed larvae showed a constant decrease from 6 DAH, except for an increase on days 13 and 14 (Fig. 2a). In these days, a high DNA content was detected in some larvae (not shown) but the cause for this is unknown. The recovery dynamics for R/D was of 1.8 R/D units day^-1^ (late-feeding) (y = 1.81x-23.181, *r*^2^ = 0.80, *p* < 0.0001) between 14 and 16 DAH, day after which values were similar to those of fed larvae. The values of R/D latency and dynamics are similar to those observed by other authors (Buckley et al. 1999). For MFS, dynamics of late-feeding larvae showed a linear decrease of 1% day^-1^ between 14 and 19 DAH (y = 19.65 - 0.95x, *r*^2^ = 0.51, *p* < 0.01). Dynamics of *D*_AZ_ of late-feeding larvae showed a slight increasing trend between 14 and 21 DAH (y =0.007 × + 0.221, *r*^2^ = 0.21, *p* < 0.05), with mean values often above those of fed larvae (Fig. 2c), although daily differences between treatments could not be established.

#### 3.2.2 Classification performance

The R/D was the only CI able to detect feeding differences in larvae ≤ 13 DAH, correctly classifying up to 69% of fed and 77% of non-fed larvae (Table 2). In larvae > 13 DAH, MFS and R/D (in this order according to their Wilks’ Lambda values) yielded the best discrimination. High discrimination values (over 78%) were achieved by R/D and MFS for fed and non-fed larvae (Table 2), whereas for late-feeding larvae these values were low (maximum of 37% by the MFS). Late-feeding larvae were confounded with fed larvae by R/D, and with any other group by MFS. In general, the *D*_AZ_ was not able to significantly ascribe the larvae to their feeding group, although a slight (high Wilks’ value) significant discrimination for fed larvae was observed only when all larvae > 13 DAH were used.

**Table 2.**
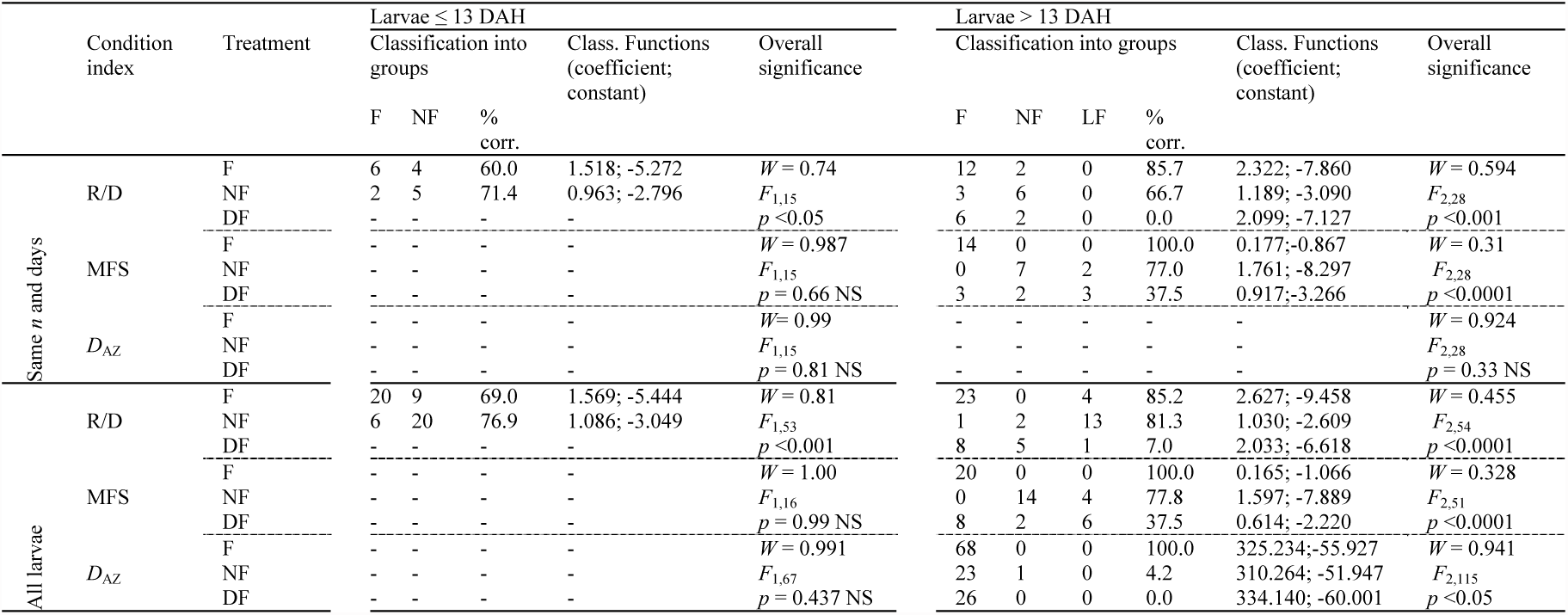
Results of the linear discriminant analysis (DA) for each condition index before (larvae ≤ 13 days after hatching, DAH) and after (larvae > 13 DAH) the total exhaustion of maternal reseves. DA was applied using 1) common sample size (*n*) and sampling days, and 2) using all the larvae. F = fed; NF = non-fed; LF = late-feeding; % corr = percent correct classification; NS = not significant; *W* = Wilks’ lambda. Significance was set after an *F* test, for which the degrees of freedom and the significance level are indicated.

#### 3.2.3 Relationship with potential muscle somatic growth and mortality

In larvae ≤ 13 DAH, no significant correlation was found between any CI and *G*_MP_ or % *G*_MP_. In larvae > 13 DAH, high R/D values were related to high *G*_MP_ (Table 3). The MFS only correlated to *G*_MP_ when all larvae > 13 DAH were used. Correlations with % *G*_MP_ were not significant for any index. The *D*_AZ_ showed no clear relationship with potential growth. Mortality rates were below 5% before 17 DAH. At 17 DAH, *M* was 9.3%, marking the initiation of the starvation-induced mass larval mortality (Fig. 2). Correlations of CIs of non-fed larvae with *M* were significant for R/D and MFS (Table 3) but could not be computed for *D*_AZ_ due to the size standardisation method. However, from Fig 2c it can be inferred that there was not a relationship between *D*_*AZ*_ and survival in the non-fed larvae.

**Table 3.**
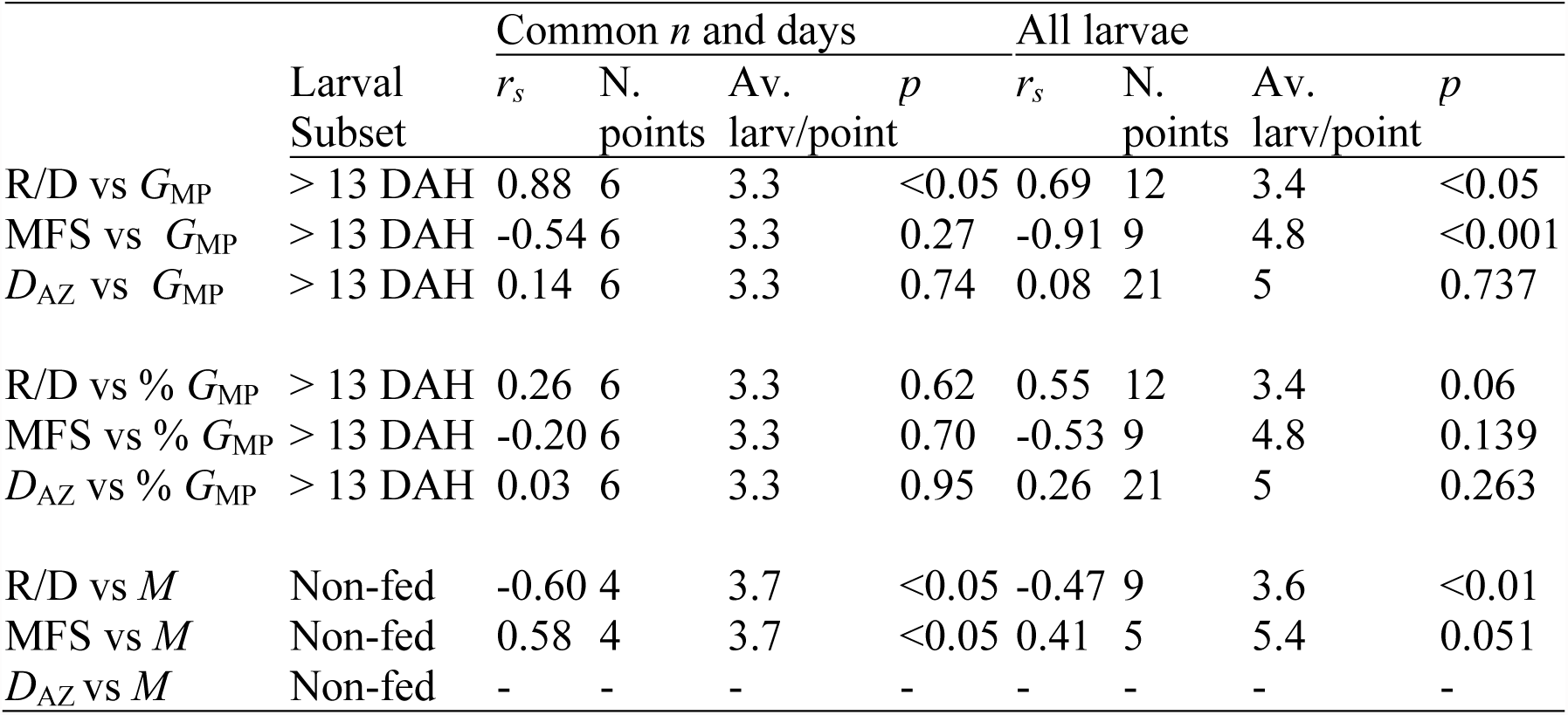
Spearman rank-correlation (*r*_*s*_) results between each condition index and daily potential muscle growth (*G*_MP_, mm^3^ day^-1^), daily percentage potential muscle growth (% *G*_MP_) and daily mortality (*M*, % day^-1^), using either a common sample size *n* and sampling dates or all larvae for the obtention of the mean values of the condition indices. For *G*_MP_ and % *G*_MP_, only larvae > 13 days after hatching (DAH) are used. For correlations with *M*, only non-fed larvae are used. N. points refers to the total data points used for correlation. Av. larv/point is the average number of larvae used for each data point.

In general, the relationships between good values of R/D and MFS and high growth rates supports the suitability of these indices for field studies in the frame of the growth-mortality hypothesis (e.g. Anderson 1988). Further, the comparable relationship of R/D and MFS with *M* suggests that these indices are not only related to growth but to starvation-induced physiological thresholds directly causing mortality. This, however, was not the case for *D*_AZ_, which did not encompass the mass mortality event (Fig. 1a). Further work on the effect of suboptimal feeding on muscle development at different time scales in a frame of physiological ecology is guaranteed.

## Acknowledgements

The authors thank C. Roldán for her help in the R/D analyses. This work was funded by the following research programmes: CYTMAR-MAR97 09-02 and Centre de Referència de Recerca i Desenvolupament en Aquïcultura (CRA-CIRIT) of the Generalitat de Catalunya. All the analyses performed comply with the Spanish laws at the time of the experiment.

